# Increased DNA methylation variability in rheumatoid arthritis discordant monozygotic twins

**DOI:** 10.1101/314963

**Authors:** Amy P. Webster, Darren Plant, Simone Ecker, Flore Zufferey, Jordana T. Bell, Andrew Feber, Dirk S. Paul, Stephan Beck, Anne Barton, Frances M.K. Williams, Jane Worthington

## Abstract

**Background:** Rheumatoid arthritis is a common autoimmune disorder influenced by both genetic and environmental factors. Epigenome-wide association studies can identify environmentally mediated epigenetic changes such as altered DNA methylation, which may also be influenced by genetic factors. To investigate possible contributions of DNA methylation to the aetiology of rheumatoid arthritis with minimum confounding genetic heterogeneity, we investigated genome-wide DNA methylation in disease discordant monozygotic twin pairs.

**Methods:** Genome-wide DNA methylation was assessed in 79 monozygotic twin pairs discordant for rheumatoid arthritis using the HumanMethylation450 BeadChip array (Illumina). Discordant twins were tested for both differential DNA methylation and methylation variability between RA and healthy twins. The methylation variability signature was then compared with methylation variants from studies of other autoimmune diseases and with an independent healthy population.

**Results:** We have identified a differentially variable DNA methylation signature, that suggests multiple stress response pathways may be involved in the aetiology of the disease. This methylation variability signature also highlighted potential epigenetic disruption of multiple RUNX3 transcription factor binding sites as being associated with disease development. Comparison with previously performed epigenome-wide association studies of rheumatoid arthritis and type 1 diabetes identified shared pathways for autoimmune disorders, suggesting that epigenetics plays a role in autoimmunity and offering the possibility of identifying new targets for intervention.

**Conclusions:** Through genome-wide analysis of DNA methylation in disease discordant monozygotic twins, we have identified a differentially variable DNA methylation signature, in the absence of differential methylation in rheumatoid arthritis. This finding supports the importance of epigenetic variability as an emerging component in autoimmune disorders.

## BACKGROUND

Low disease concordance rates between monozygotic (MZ) twins (~15%) have revealed that environmental exposures are important in rheumatoid arthritis (RA) [1]. Many putative environmental risk factors have been investigated, including exposure to cigarette smoke, hormone influences, infection, vitamin D intake and dietary factors [2, 3], but few have been robustly confirmed.

Epigenetics is the study of heritable modifications of DNA which can alter gene expression without changing the DNA sequence, and which can be influenced by environmental factors, such as smoking [4]. The most widely studied epigenetic phenomenon is DNA methylation, which may act as a composite measure of numerous environmental exposures, making it an intriguing candidate for investigation of diseases that involve both genetic and environmental factors, such as RA.

Current evidence suggests that DNA methylation changes are associated with RA [5–10] but whether this is due to intrinsic genetic differences, which can also influence DNA methylation, is not yet known. Disease-discordant MZ twin pairs offer an ideal study design as they are matched for many factors, including genetic variation and as such they offer a crucial advantage in epigenetic studies [11]. Differences in methylation between MZ twins may capture the effects of environmentally driven mechanisms, independent of genetically driven changes. Two small epigenome-wide association studies (EWAS) of DNA methylation in MZ twins discordant for RA have reported conflicting results. The first (n=5 pairs) identifying no significant changes associated with RA using the GoldenGate assay [12], while the second (n=7 pairs) identified no significant differentially methylated positions (DMPs), but one significantly differentially methylated region (DMR) using the CHARM platform [13]. Due to the small sample sizes of both studies, they were underpowered to detect subtle methylation differences [14] and have limited scope to characterise the epigenomic landscape of RA discordant twins.

To our knowledge, all studies of DNA methylation in relation to RA have focussed on the identification of DMPs or DMRs, which describe differential DNA methylation levels at a particular CpG site or closely spaced group of CpG sites, respectively. In DMPs and DMRs, one group has a consistently higher level of DNA methylation than the comparison group (eg, when comparing RA patients with healthy controls). Differentially variable positions (DVPs) are another type of epigenetic variation, the importance of which has recently been elucidated in type 1 diabetes (T1D), cervical and breast cancer [15–18]. DVPs are CpG sites that do not necessarily have a large difference in mean DNA methylation, therefore may not be classed as DMPs; however, they have substantial difference in the range of DNA methylation values between comparison groups.

We have examined genome-wide DNA methylation in both a DMP and DVP context using the Infinium HumanMethylation450 BeadChip array (Illumina) in whole blood from 79 MZ twin pairs discordant for RA from two independent cohorts (Manchester and TwinsUK, see Figure 1). We identified a DVP signature in the absence of a DMP signature, that suggest potential roles for multiple stress response pathways and potential epigenetic disruption of RUNX3 transcription factor binding sites in RA aetiology. We also identified shared DVPs in both RA and T1D, indicating potential shared pathways for autoimmune disorders.

**Figure 1.**
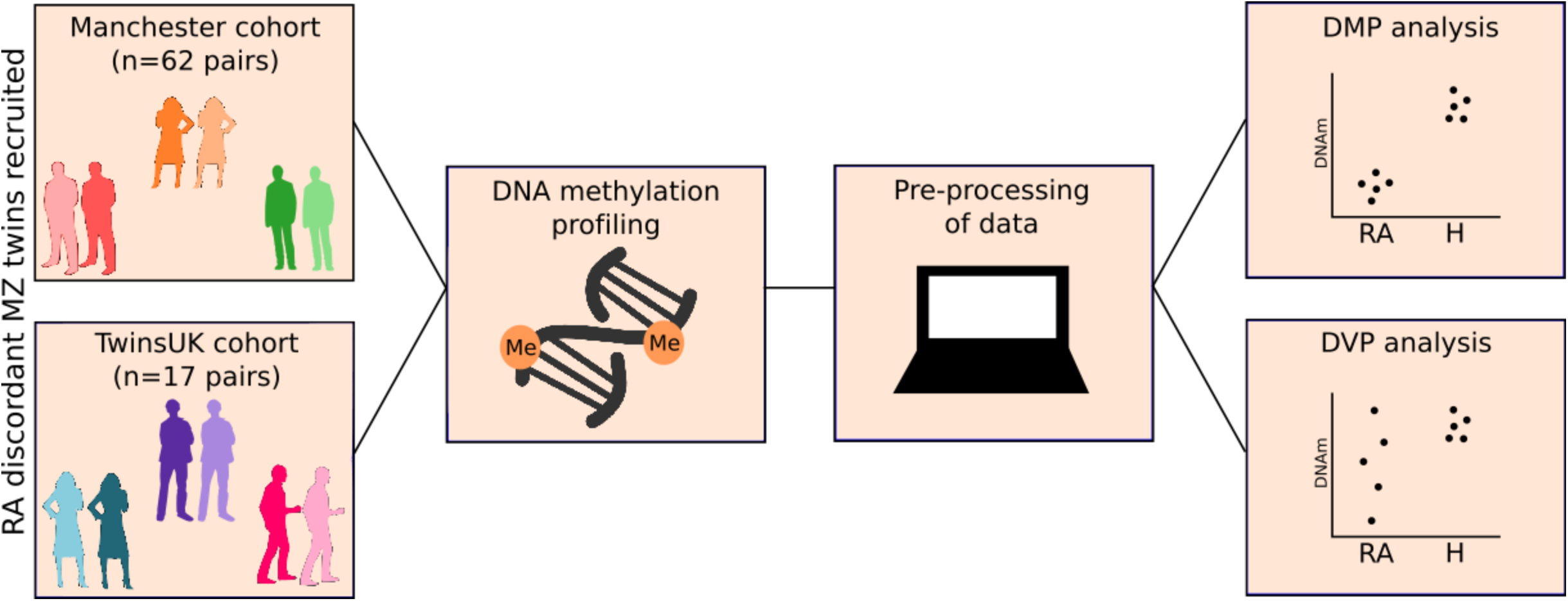
Overview of study design. Rheumatoid arthritis discordant twin pairs were recruited from the RA twins study in Manchester and TwinsUK in London, and genome-wide DNA methylation was investigated in the context of both differentially methylated positions and differentially variable positions.

## PATIENTS & METHODS

### TwinsUK participants

Twin pairs discordant for RA were identified from the TwinsUK register [19]. RA status was assessed through questionnaires between 1997 and 2002. In addition, an advertisement was published in the National Rheumatoid Arthritis Society newsletter in spring 2013 to recruit twin volunteers with RA. All MZ twins who answered positively were phone-interviewed by a rheumatology clinical fellow to confirm the diagnosis of RA based on the American College of Rheumatology 1987 criteria (n=17 RA twins). In case of unclear diagnosis of RA, participants were reviewed in clinic, or were excluded. In addition, all patients willing to attend a visit were examined clinically (n=11 RA twins) by a clinical fellow under the supervision of a consultant rheumatologist. Visits included detailed medical history, review of symptoms, past and present medication (NSAIDS, DMARDS and/or biological agents) as well as joint examination. Blood samples were collected from all subjects, from which DNA and serum were extracted and stored at -80 °C.

The healthy co-twins were also reviewed at the clinical visit. Non-RA status was supported by both clinical and immunological details, as all non-RA twins were seronegative, except one who was rheumatoid factor (RF) positive but clinically unaffected.

### Manchester participants

Patients were selected from the Nationwide Rheumatoid Arthritis Twin Study based at the University of Manchester [1]. Twins were recruited in 1989 using a dual strategy: 1) all UK rheumatologists were contacted and requested to ask all of their patients with RA whether they were a twin; 2) a multimedia campaign was targeted to patients in whom RA had been diagnosed and who had a living twin. Both members of each twin pair were visited at home by trained research nurses who recorded each subject’s detailed medical history and demographic characteristics and performed joint examinations. Blood samples were collected from all subjects, from which DNA and serum were extracted and stored at -80 °C.

### Measurement of genome-wide DNA methylation

For each sample, 500ng DNA was bisulfite-converted using EZ DNA methylation kits (ZYMO Research) according to the manufacturer’s amended protocol for use with the Infinium HumanMethylation450 BeadChip (Illumina). Epigenome-wide methylation was assessed using the Infinium HumanMethylation450 Assay (Illumina) and the BeadChips were then imaged using the Illumina iScan System.

### Quality control and pre-processing of HumanMethylation450 data

All data analysis was performed in R 3.4.1 (R Development Core Team) using the minfi [20], ChAMP [21] and CpGassoc packages [22]. Data quality for each sample was assessed by visual inspection of kernel density plots of methylation beta-values and by comparing median log2 intensities recorded in both the methylated and unmethylated channels. Probes which failed a detection-P value of 0.01, probes mapping to the sex chromosomes, probes containing a SNP within two base pairs of the measured CpG site, cross reactive probes (according to Norlund, 2013) and probes with a bead count of <3 in at least 5% of samples were removed prior to analysis. Raw beta values were logit transformed to M-values following subset-quantile within array normalisation (SWAN) and principal component analysis (PCA) was performed to capture any potential technical variation. Distinct cell populations are known to have different DNA methylation signatures [23]. Therefore, to assess if cell composition differs between healthy and RA affected twins, and whether this may confound downstream analysis, we estimated cell composition for each sample using the reference-based Houseman method to infer relative proportions of cells [24]. Differences in cell composition between groups was tested using a Welch two sample t-test.

### Identification of differentially methylated positions (DMPs)

A mixed effects model was used to test for DMPs from beta values using the CpGassoc package, adjusting for sibling-pair effects as a random covariate. Factors associated with the first 4 principal components (PCs) were included in the model as fixed covariates. False discovery rate was calculated using the Benjamini and Hochberg method [25], and a significance threshold of 0.05 was used. Power to detect differential DNA methylation was estimated using the calculations presented in [14], with genome-wide significance threshold set to 1E-06 and the false discovery rate controlled at 0.05.

### Identification of differentially variable positions (DVPs)

Differential DNA methylation variability was tested in the current study using the recently developed iEVORA algorithm [16], which employs a modified version of the Bartlett’s test to test for differences in variability, in combination with a standard t-test to subsequently rank the identified DVPs. A significance q-value threshold of 0.001 was applied for the differential variability test, while a significance p-value threshold of 0.05 was applied for the differential means.

### Assessment of DVP signature in an independent healthy population

In order to assess if the DVP signature identified between RA discordant twins was present in an independent healthy cohort, methylation variability was assessed in the BIOS cohort described in [26]. Briefly, this dataset consisted of HumanMethylation450 profiles generated from three Dutch cohorts, from which 156 profiles were randomly selected to test methylation variability at the DVP sites. The variance and range of methylation values were calculated for each CpG site in the DVP signature, stratified by directionality of variability in the signature (ie, if DVPs were hypervariable in healthy or RA twins).

### Feature enrichment analysis

To investigate if DVPs identified in RA discordant twins were enriched in particular CpG island associated features, or in certain gene features, an enrichment analysis was performed. All CpG sites included in analysis were annotated using the HumanMethylation450 manifest. Repeated random sampling (n=1000) of all probes that passed quality control was used to assess enrichment of features associated with DVPs [27].

### Pathway Analysis

Pathway analysis was performed within the MissMethyl package [28] using the gometh function. Methylation arrays have a significant bias in pathway analysis due to the differential distribution of probes across different genes [29]. For example, on the HumanMethylation450 BeadChip (Illumina), the number of probes on each gene represented on the array ranges from 1 to 1299. Consequently, during standard pathway analyses, genes with a large number of probes present on the array are more likely to be implicated in significant pathways. The MissMethyl package adjusts for such bias using a modified hypergeometric test to test for over-representation of the selected genes in each gene set. Pathways were ranked by p-value for over-representation of the gene ontology terms (p<0.05). False discovery rate (FDR) correction was not applied because biological pathways are not independent from each other, and the FDR procedure is only valid when tests are independent [25]. Additionally, we performed Gene Set Enrichment Analysis [30] on DVP-associated genes to corroborate top-ranked pathways.

### Overlap analyses

Using a meta-analysis approach, the DVPs from the current study were compared with DVPs and DMPs identified in previously performed large-scale EWAS of various autoimmune disorders. Studies were selected that had performed a site-specific genome-wide study of DNA methylation (eg, using methylation microarrays) in an autoimmune disease, with at least 100 individuals included in the study. The two qualifying studies focussed on T1D [15] in a discordant MZ twin approach, and RA [7] in an unrelated case-control approach. Lists of statistically significant DVPs (q<0.001) and DMPs (Bonferroni corrected p<0.05) respectively reported in each study were overlapped with DVPs identified in the current study. This allowed identification of DVPs which were common across different diseases and different study designs.

## RESULTS

### Patient characteristics

DNA samples from 79 MZ twin pairs discordant for RA were available from the Nationwide Rheumatoid Arthritis Twin Study (n=62 twin pairs) and from the TwinsUK cohort (n=17 twin pairs). Patient characteristics are summarised in Table 1. There was no significant difference regarding smoking status between RA and non-RA co-twins (p=0.53). Of the RA co-twins in the study, 59% were seropositive (anti-CCP and/or RF), whereas 9% of the non-RA twins were RF positive. Of the RA co-twins, 52% were taking disease-modifying anti-rheumatic drugs (DMARDs) at the time of the blood sampling, with the most prescribed being methotrexate (n=9). Other commonly prescribed mono- or bi-therapy DMARDs included penicillamine (n=8), gold (n=8), sulphasalazine (n=7) and hydroxychloroquine (n=5), reflecting the prescribing practices at the time when the data was collected for the larger group of twins. Treatment with DMARDs was not associated with any of the top 20 principal components, and during assessment with multidimensional scaling of the top 1000 most variable probes, treatment with DMARDs did not separate out different groups (Supplementary Figure 1), therefore it was not adjusted for in the analysis. This study had >80% power to detect a mean methylation difference of 4% and 13% between the RA and non-RA twins, at the 5% and genome-wide significance threshold, respectively.

**Table 1.**
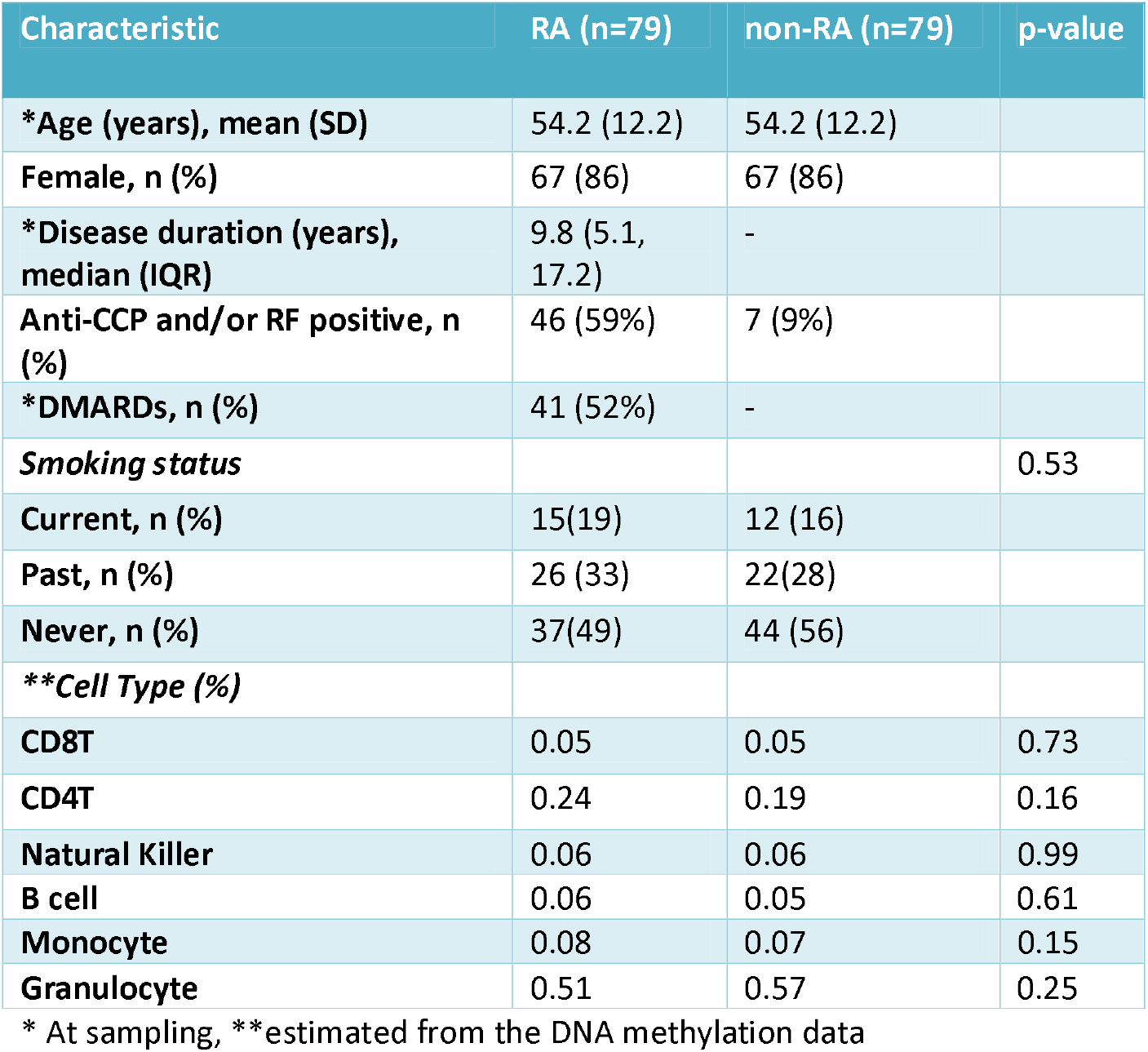
Characteristics of the RA discordant twin pairs

### DMP analysis

Following stringent probe filtering, 430,780 probes were available for further analyses in the dataset. Potential confounding factors including gender, age, smoking habits, cell composition, cohort, position on array and BeadChip ID were all included as covariates in the linear regression. None of the probes investigated were significantly differentially methylated between the RA and non-RA twins following correction for multiple testing, using a false discovery rate threshold of 0.05. The mean difference in methylation between RA discordant twins for the probes with the smallest adjusted p-values (p>0.13) was less than 4% (Table 2). To assess the influence of differences in cell type composition on the epigenetic profiles, we inferred differential cell type proportions based on the DNA methylation data [24]. Proportions of each cell type were compared between the two comparison groups, and there were no significant differences (p>0.15) in cellular composition between RA and healthy co-twins (Table 1, Supplementary Figure 2). Furthermore, adjustment for cell composition during DMP detection did not affect the results qualitatively.

**Table 2.** Top 20 differentially methylated positions between RA twins and non-RA twins. Probe names are shown, along with methylation levels, intra-pair methylation difference, unadjusted p-value and probe annotation.

| Probe | Non-RA beta | RA beta | Diff | P-value | Chr | Relationship to gene | Gene symbol |
| --- | --- | --- | --- | --- | --- | --- | --- |
| **cg26547058** | 0.763 | 0.800 | 0.037 | 4.89E-07 | 8 |  |  |
| **cg07693617** | 0.758 | 0.788 | 0.030 | 5.15E-07 | 1 | Body | PRKCZ |
| **cg07636225** | 0.849 | 0.867 | 0.018 | 9.33E-07 | 11 | Body | RTN3 |
| **cg03517226** | 0.709 | 0.716 | 0.007 | 1.18E-06 | 16 | 5'UTR | ANKRD11 |
| cg20666386 | 0.184 | 0.180 | -0.004 | 1.89E-06 | 11 | 1stExon | DGKZ |
| cg17501210 | 0.766 | 0.740 | -0.026 | 2.54E-06 | 6 | Body | RPS6KA2 |
| cg26964117 | 0.775 | 0.796 | 0.021 | 2.56E-06 | 19 | Body | PIH1D1 |
| cg26701826 | 0.396 | 0.419 | 0.023 | 2.71E-06 | 4 | 5'UTR | SGMS2 |
| cg06040872 | 0.810 | 0.822 | 0.012 | 2.92E-06 | 17 | Body | CCL18 |
| cg25487804 | 0.867 | 0.876 | 0.009 | 3.38E-06 | 11 | TSS1500 | OSBPL5 |
| cg06128521 | 0.900 | 0.905 | 0.005 | 3.56E-06 | 17 |  |  |
| cg16640599 | 0.421 | 0.447 | 0.026 | 3.58E-06 | 4 | Body | SEC24D |
| cg02445229 | 0.719 | 0.737 | 0.018 | 3.85E-06 | 19 |  |  |
| cg01769457 | 0.039 | 0.043 | 0.004 | 3.90E-06 | 1 |  |  |
| cg15293582 | 0.704 | 0.703 | -0.001 | 4.06E-06 | 10 | TSS1500 | PRF1 |
| cg07090025 | 0.091 | 0.100 | 0.009 | 4.21E-06 | 1 | TSS200 | SSU72 |
| cg06791979 | 0.795 | 0.799 | 0.004 | 4.44E-06 | 11 | Body | RTN3 |
| cg01934296 | 0.939 | 0.931 | -0.008 | 4.45E-06 | 1 | Body |  |
| cg26272088 | 0.819 | 0.838 | 0.019 | 4.54E-06 | 19 | Body | IRGC |
| cg18460107 | 0.674 | 0.679 | 0.005 | 4.71E-06 | 17 | TSS200 | SEPT9 |

Chr=chromosome

### Rheumatoid arthritis associated DVPs

Variability of DNA methylation has been implicated in T1D, cervical and breast cancer [15–17]. We used the recently developed iEVORA algorithm [16] to test if DNA methylation variability was significantly associated with RA status between disease discordant MZ twins. In a group-wise test for differential variability between RA discordant MZ twins, 1171 DVPs were identified at a stringent false discovery rate of <0.001. An example of the six top-ranked DVPs is shown in Figure 2 and the annotation of the top 20 DVPs is summarised in Table 3 (full list of DVPs provided in Supplementary Table 1). These DVPs were enriched in CpG sites that did not map to CpG Islands, and were enriched in the body and 3’UTR of genes (Supplementary Figure 3).

**Figure 2.**
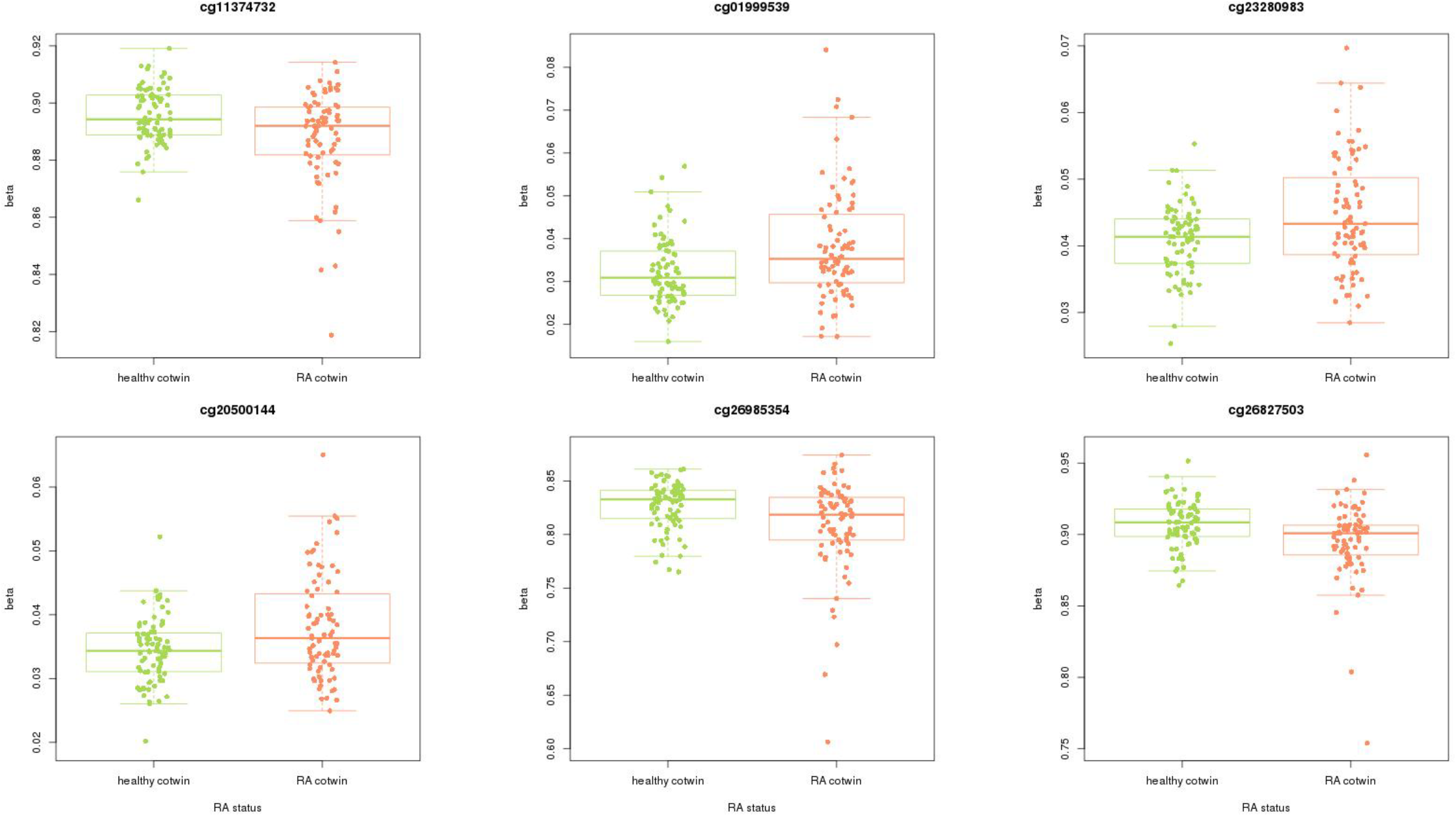
CpG plots for six top ranked differentially variable positions in RA discordant MZ twins. Cpg sites shown are cg11374732 (Bartletts test p-value=4.09E-06), cg01999539, cg23280983, cg20500144, cg26985354, cg26827503. Hypervariability of differentially variable positions was enriched in RA twins. Boxplots indicating the mean methylation and range of methylation values are shown overlaid with scatterplots indicating DNA methylation measurements of individual samples.

**Table 3.**
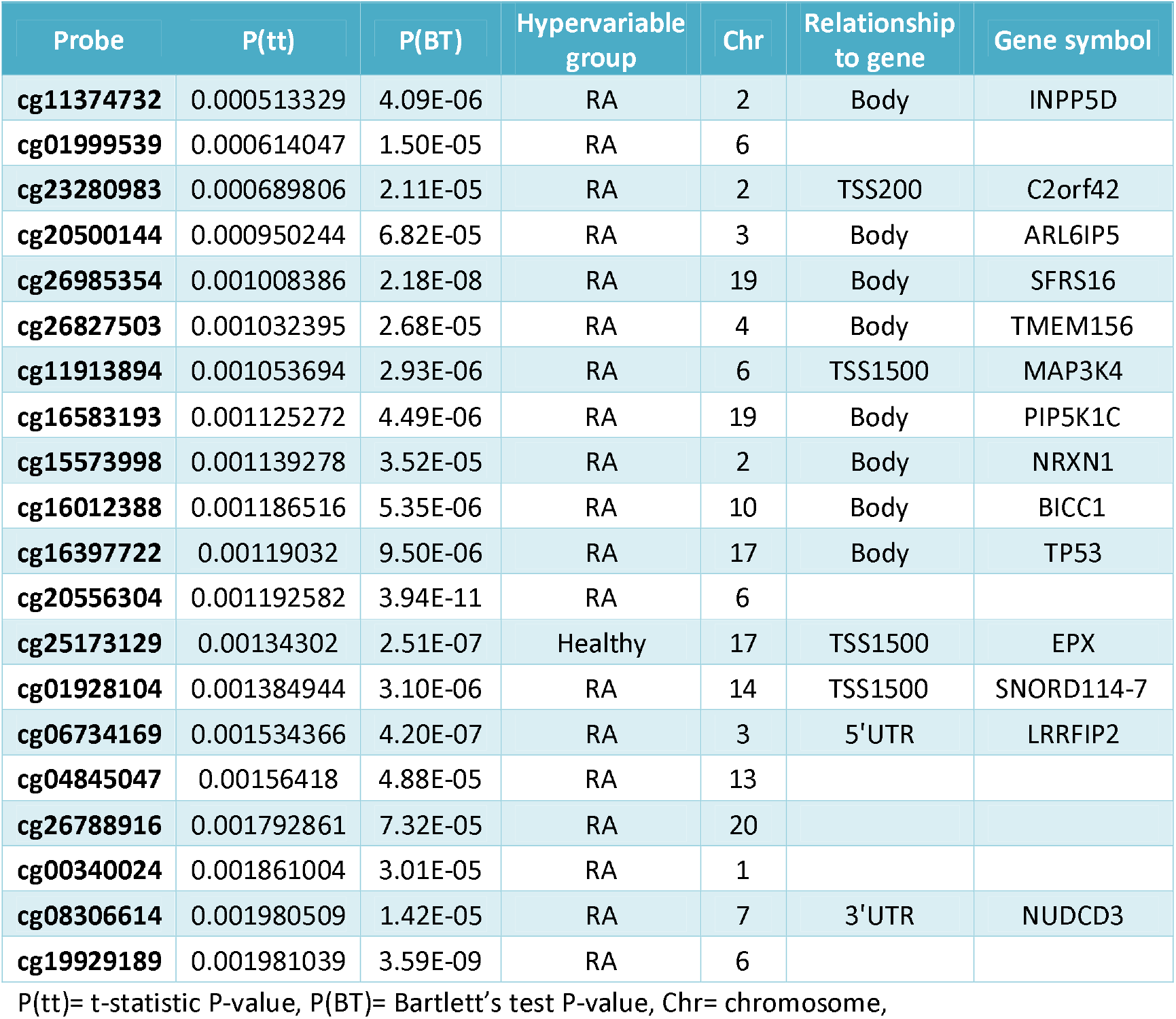
Top 20 differentially variable positions between RA affected and non-RA twins. Probe names are shown, along with t-statistic p-value, Bartlett’s test for differential variability, which group was hypervariable, and probe annotation.

Of the 1171 DVPs, 763 were hypervariable in the RA twins, indicating an enrichment of methylation variability in disease-affected individuals. DVPs that were hypervariable in RA twins were enriched in the 3’UTR of genes and gene bodies, while DVPs that were hypervariable in healthy twins were enriched in gene bodies (Figure 3). The underrepresentation of DVPs in CpG Islands, particularly in DVPs that were hypervariable in healthy twins, indicates that these regions are more epigenetically stable.

**Figure 3.**
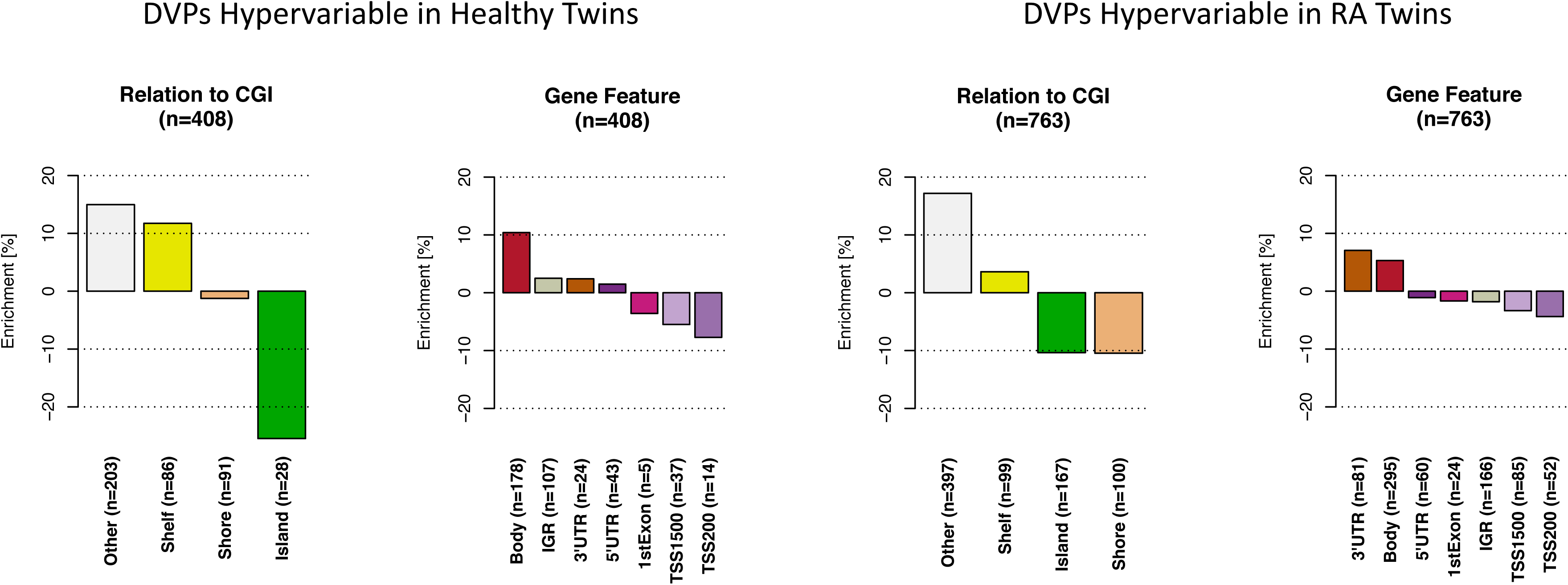
Feature enrichment for RA associated differentially variable positions which are hypervariable in healthy and RA affected twins respectively. Differentially variable positions which are hypervariable in the healthy twins were enriched in CpG Island shelves and non-CpG island associated regions, and in the bodies of genes. Meanwhile differentially variable positions enriched in RA affected twins were enriched in 3’UTRs of genes, gene bodies and non-CpG Island associated regions.

To investigate the DNA methylation variability of RA-associated genes, DVPs were annotated and the gene associated with each DVP was compared to RA-associated genes identified during genetic studies. Meta-analysis of RA susceptibility loci has previously identified 98 genes associated with 101 genetic variants [31]. Of these 98 RA-associated genes, five contained at least one DVP (*CLNK, JAZF1, ICOSLG, NFKBIE* and *BLK*). *JAZF1* contained two DVPs, both of which map to the body of the gene. Further, when the 377 genes with nominal association to RA from the same study were investigated, 15 genes contained at least one DVP. The presence of genetic and epigenetic variants in the same susceptibility genes indicates that disease-related changes in gene function or expression could be implemented by different mechanisms.

Functional annotation of the top ranked DVPs (ranked by T statistic P-value) showed that the second and third most highly ranked CpG sites (cg01999539 and cg23280983) overlap with the binding site of the transcription factor RUNX3. This differential variability of DNA methylation could potentially be influencing the binding of this transcription factor in multiple locations throughout the genome. Several studies have implicated RUNX3 in the development of immune related diseases including Crohn’s disease, ankylosing spondylitis, psoriasis and ulcerative colitis (reviewed in [32]), and SNPs which disrupt RUNX binding sites have also been associated with RA [31, 33]. The disruption of the expression of RUNX3 transcription factors has also been found to alter the suppressive function of regulatory T cell in human cells and in mice, suggesting a potential functional consequence of the methylation changes observed in RUNX3 binding sites which warrants further investigation in RA [34].

### Pathway analyses of rheumatoid arthritis associated DVPs

Pathway analyses identified an enrichment of the RA-associated DVPs in pathways (p<0.05) involved in response to cellular stress (Supplementary Table 2). A pathway involving ubiquitination of the protein K63 was also identified as enriched; this pathway has been found to modulate oxidative stress response [35] which has a role in RA pathogenesis [36]. When the pathway analysis was restricted to DVPs which are hypervariable in non-RA twins, there was an enrichment for immune-related processes in the top ranked pathways (Supplementary Table 3). Two of the five top-ranked pathways were related to T cell cytokine production, a critical process in the development of RA.

### Overlap with previously identified rheumatoid arthritis associated DMPs

Changes in DNA methylation have previously been associated with RA, however, these studies were performed in unrelated case-control study designs. We hypothesised that the DMPs, which are found to be consistently differentially methylated across comparison groups in the analysis of unrelated individuals, may overlap with the differentially variable methylation signature identified in the current study. To test this, we overlapped the DMPs from the largest EWAS of RA to date with the DVPs identified in the current study, to assess if alternative study designs and approaches had identified common epigenetic variants.

The previous study was performed in 691 unrelated individuals (n=354 RA cases and 337 unrelated controls) and identified 51,476 DMPs which were putatively associated with RA, achieving a P<0.05 following Bonferroni correction [7]. Of the 1171 DVPs identified in the current study, 132 overlapped with DMPs identified in the unrelated RA EWAS. Of these, 123 DVPs were hypervariable in the RA-affected twins in the current study. In pathway analysis of these overlapping epigenetic variants, one of the top ranked pathways was associated with low-density lipoprotein receptor activity (p=0.003), which associates closely with the protein produced by the *LRPAP1* gene, the methylation of which was recently reported as a potential biomarker of anti-TNF treatment response in RA patients [37].

### Overlap with type 1 diabetes associated DVPs

Genetic susceptibility loci identified in RA have also been found to confer risk for other autoimmune disorders. RA and T1D are both common autoimmune diseases with many characteristics in common, and it is possible that similarities in epigenetic profile between the two disorders may provide insight to the development of autoimmune diseases in general. To test whether epigenetic variants have commonality across different autoimmune disorders, we tested for overlap between the RA associated DVPs identified in the current study, and a set of DVPs recently identified in a T1D twin study.

A recent study of DNA methylation in T1D discordant monozygotic twins (n=52 twin pairs) identified 16915 unique DVPs that were hypervariable in T1D across three cell types. Overlap analysis of the genes associated with these probes identified 496 genes that are associated with both RA-DVPs and T1D-DVPs, which is more than expected by chance (p=1.6e-29), measured using the phyper test for hypergeometric distribution. An overlap analysis of methylation probe IDs identified 69 specific probes that overlap between RA-associated DVPs and T1D-associated DVPs. Permutation testing with 10,000 iterations indicated this overlap is more than expected by chance (p=0.001). Pathway analysis of these probes identified many immune-related pathways including a pathway involved in NF-KB cascade modulation, a process that is important in inflammation and immune responses. Many of the top-ranked pathways identified were related to embryonic development, including neural, embryonic and epithelial tube formation. The overlapping probes were enriched in 3’UTR and intergenic regions (Supplementary Figure 4).

### Assessment of methylation variability signature in an independent healthy cohort

The DVP signature identified in RA discordant twins was tested in an independent cohort of healthy individuals from the BIOS cohort [26] to assess if the methylation variability signature was specific to RA or can also be detected in the general population. Variance and range statistics were generated for each DVP for three comparison groups (RA co-twins, healthy co-twins and healthy BIOS individuals). The aggregated variance and range statistics for the sites were then plotted to compare distribution across the three groups, split by direction of variability. The DVPs which were hypervariable (n=763) in RA affected twins had a lower variance and a lower range of methylation values in both healthy twins and the BIOS healthy cohort when compared with RA affected twins (Figure 4). The DVPs that were hypervariable (n=408) in the healthy co-twins were also found to have the same trend when compared with the BIOS cohort, with larger variance and range in the healthy co-twin and BIOS groups than in the RA affected group (Supplementary Figure 5).

**Figure 4.**
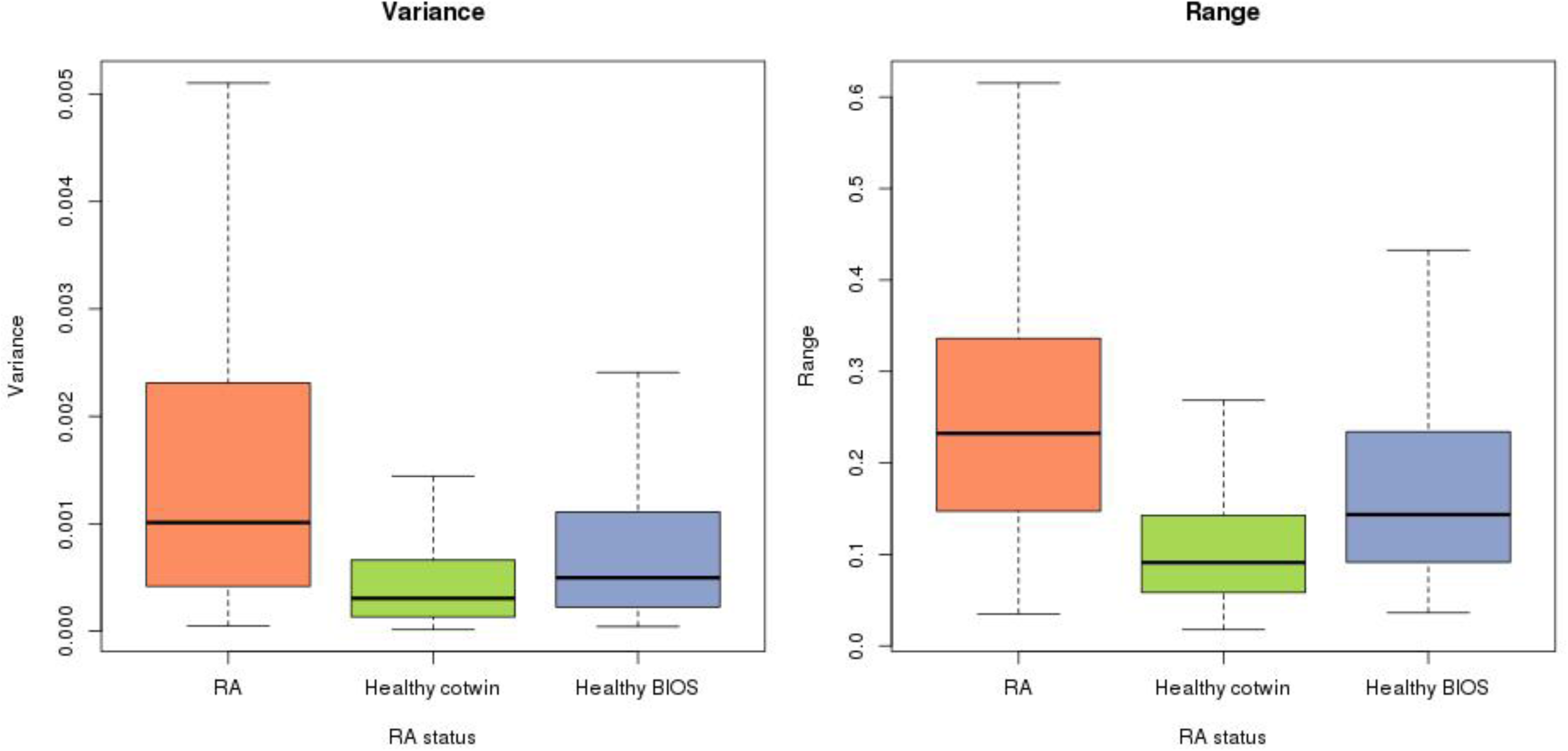
Variance and range for DVPs which were hypervariable in RA co-twins. Variance and range were calculated for each of the 763 DVPs found to be hypervariable in RA co-twins, plotted for three comparison groups; RA co-twins, healthy co-twins and an independent cohort of healthy individuals.

## DISCUSSION

In the largest study of DNA methylation in RA-discordant MZ twins performed to date, we have identified a significant differential variability signature in RA. Differentially methylated positions were not identified following adjustment for multiple testing in 79 pairs of disease discordant twins. The identification of a differentially variable signature in the absence of a differentially methylated signature supports the recent findings of an EWAS of T1D discordant monozygotic twins, which identified 10,548 DVPs in B cells, 4314 in T cells and 6508 in monocytes [15]. While the T1D study had a smaller sample size (n=52 T1D discordant twin pairs), it had increased power to detect methylation differences due to the use of individual cell types. A limitation of the current study is that it was performed in whole blood, making it more difficult to identify subtle methylation differences. However, cell estimates were imputed from the methylation data using a reference-based statistical algorithm [24] indicating that there were no significant differences in proportions of cells tested. The overlap of the RA associated DVPs with T1D associated DVPs identified in individual cell types is interesting as it indicates that at least a subset of DVPs identified in individual cell types can also be identified using whole blood. Another limitation of the study is that the samples were sourced from two cohorts, which inevitably confers a batch effect in the data. However, as each RA-affected individual is matched with their unaffected co-twin from the same study, the effect of this on the analysis is negligible. While the biological implications of DVPs are not yet fully understood, such DVPs have been found to be temporally stable over five years in T1D [15]. Further longitudinal studies are required to assess if this is the case in RA. It is also important to consider that the methylation variability detected may reflect either cause or consequence of the disease, which warrants further investigation into the temporal origins and functional consequences of this methylation variability signature.

RA associated DVPs were enriched in pathways involved in the regulation of response to stress, including stress-activated kinase signalling cascades. While these pathway associations are based on bioinformatics analysis of methylation data, they have intriguing links to inflammatory pathways which warrant further functional investigation in RA. For example, stress kinases have previously been found to be induced by pro-inflammatory cytokines in RA, and the stress-activated protein kinase pathway has been shown to be active in RA synovium, while not being active in the synovium of patients with osteoarthritis [38]. These findings indicate that the inflammatory component of RA could potentially induce the observed variability of DNA methylation in stress response pathways. The variability of DNA methylation in stress-response regulating pathways might reflect the adaptation of these cells to stress-inducing conditions which are present in RA, such as increased levels of cytokines and induction of oxidative stress. These findings have lead us to propose a new working model of the development of RA (Figure 5) illustrating the potential role of DNA methylation variability and stress response pathways in the aetiology of disease.

**Figure 5.**
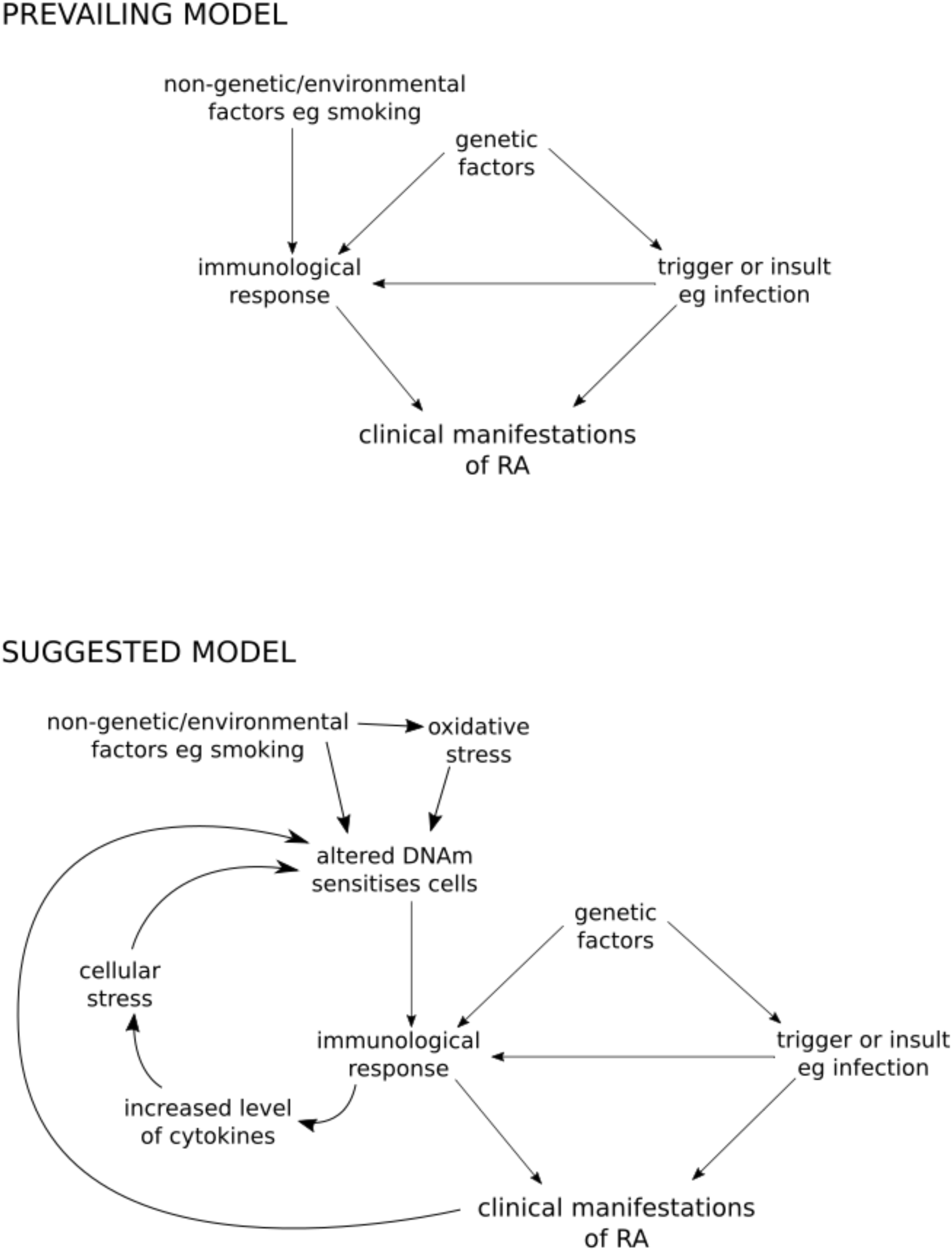
Prevailing and suggested model of RA disease development. The current model of RA aetiology primarily involves the contribution of genetic and non-genetic factors including triggers such as infection, which mediate an immune response and ultimately contribute to the clinical manifestation of RA. Our suggested model of RA development incorporates the potential role of DNA methylation alongside the contributing factors of oxidative and cellular stress.

The RA associated DVPs were also enriched in a pathway controlling K63 protein ubiquitination. The ubiquitination of K63 acts as a modulator of oxidative stress response, which induces thioredoxin, a catalyst found to be overexpressed in RA patients [39]. Thioredoxin has also been found to activate the NF-KB pathway [40], which we observed to be enriched with DVPs in both RA and T1D affected twins.

Disease-associated epigenetic alterations have been hypothesised to be caused by chronic cellular stress, which can be induced by inflammation [41]. Accumulating evidence indicates that these stress induced changes in the epigenetic landscape cause changes in cellular state and function, which can contribute to disease development. The enrichment of RA-associated DVPs in stress response related pathways supports this hypothesis, and suggests that as well as altering cellular state, such epigenetic changes are also modulating the response to cellular and oxidative stress. This could be perpetuating the disease phenotype by repressing cellular stress response mechanisms in cells exposed to inflammation.

Meta-analysis of two other autoimmune EWAS studies identified a set of 496 genes that contain DVPs in both RA and T1D. This provides an intriguing possibility of common pathways in which DNA methylation is hyper-variable in autoimmune disorders. These may provide novel pathways for treatment, and generate hypotheses regarding autoimmune disease pathogenesis. For example, one of the overlapping genes was *PRKCZ*, which was also the gene containing the second most differentially methylated probe in the current study. This gene was previously found to be hypermethylated in RA fibroblast-like synoviocyte cells [42], and has also been found to be hypomethylated in T1D monocytes and whole blood [43]. The largest RA EWAS of unrelated individuals identified 132 DMPs which were found to overlap with the DVPs identified in the current study. Pathway analysis of these sites identified a pathway associated with the *LRPAP1* gene, which was recently identified as a biomarker of treatment response in RA [37]. This overlapping signature of differential variability on methylation in two autoimmune diseases suggests that epigenetics plays a role in autoimmunity, which warrants further investigation in functional studies to elucidate its’ role in autoimmune disease pathogenesis.

## CONCLUSIONS

In a genome-wide investigation of DNA methylation in RA-discordant MZ twins, we have identified differential variability of DNA methylation, but no statistically significant DMPs. This supports the findings of a recent investigation of DNA methylation in T1D discordant monozygotic twins, which identified a disease associated DVP signature in the absence of a substantial DMP signature. Due to the influence of genetic components in establishing DNA methylation [44], our study indicates that differentially methylated positions that have previously been associated with RA [6–10] do not replicate in cohorts of disease-discordant monozygotic twins, which are less confounded by genetic heterogeneity. Furthermore, we identified a series of stress-response associated pathways which may potentially play a role in RA aetiology. These pathways interact with pro-inflammatory cytokines known to be integral in the development of RA, thus are of direct relevance to RA pathogenesis and could provide potential targets for RA therapy development. The role of stress response in RA pathology warrants further investigation to determine the downstream functional effects of this DNA methylation variability, and to further characterise the role of variability of DNA methylation in complex diseases such as RA.

## Acknowledgements

The authors thank the UCL Medical Genomics group, particularly Ismail Moghul, for their support, Andrew Teschendorff for useful conversations, and Paul Guilhamon for sharing his feature enrichment analysis method. The authors also thank Antonino Zito and Kerrin Small from TwinsUK for providing data for analysis. The authors also thank Christopher Bell and Richard Acton for their contribution to the analysis.

## LIST OF ABBREVIATIONS

MZ: Monozygotic
RA: Rheumatoid arthritis
DMPs: Differentially methylated positions
DMR: Differentially methylated region
DVPs: Differentially variable positions
T1D: Type 1 diabetes
RF: Rheumatoid factor
SWAN: Subset-quantile within array normalization
PCA: Principal component analysis
PCs: Principal components
FDR: False discovery rate
EWAS: Epigenome-wide association study
DMARDs: Disease-modifying anti-rheumatic drugs
NF-KB: Nuclear factor kappa-light-chain-enhancer of activated B cells
UTR: Untranslated region

## DECLARATIONS

### Ethics approval and consent to participate

The study was approved by the North-West Haydock Ethics Committee (MREC 99/8/84) and the St Thomas’ Hospital Ethics Committee.

### Authors’ contributions

The study was conceived and designed by APW, JW and FMW. Sample collection was performed by FZ, AB, FMW and JW. Experimental work was performed by APW. Statistical analysis of was performed by APW and SE with support from DP, JTB, AF, DSP and SB. The manuscript was written by APW with support from DP, SE, FZ, JTB, AF, DSP, SB, AB, FMW and JW. All authors read and approved the manuscript.

### Availability of data and material

The HumanMethylation450 data will be made available in the ArrayExpress Archive of Functional Genomics Data (European Bioinformatics Institute).

### Consent for publication

Not applicable.

---

Supplementary Figure 1. **Multidimensional scaling plot of DMARD use in RA discordant twins.** MDS plot showing similarities and differences between samples using Euclidian distances based on methylation values of the 1000 most variable positions in analysis. The samples are coloured by DMARD use and the treatment does not differentiate samples into discrete groups.

Supplementary Figure 2. **Cell composition estimates for RA discordant twins.** Plot shows relative proportions of each cell type for RA and non-RA co-twins, estimated from DNA methylation data using bioinformatics methods. Proportions of each cell type were not found to be statistically different between the two comparison groups (p=0.05).

Supplementary Figure 3. **Feature enrichment for differentially variable positions.** Differentially variable positions are primarily enriched in regions not associated with CpG Islands. Gene feature enrichment showed that differentially variable positions were enriched in gene bodies and 3’UTR regions of genes.

Supplementary Figure 4. **Feature enrichment for DVPs identified in both RA and type 1 diabetes disease-discordant twins.** DVPs were not enriched in CpG Island associated features (left), but were enriched in 3’UTR sites of genes.

Supplementary Figure 5. **Variance and range for DVPs which were hypervariable in healthy co-twins.** Variance and range were calculated for each of the 408 DVPs found to be hypervariable in healthy co-twins, plotted for three comparison groups; RA co-twins, healthy co-twins and an independent cohort of healthy individuals.

Supplementary Table 1. Full list of differentially variable positions (n=1171) between RA affected and non-RA twins. Probe names are shown, along with t-statistic p-value, Bartlett’s test for differential variability, which group was hypervariable, and probe annotation.

Supplementary Table 2. Pathways enriched in differentially variable positions identified in RA discordant twins. Pathway analysis was performed using the gometh function in the MissMethyl package. Pathways are ranked by P-value (P<0.05).

Supplementary Table 3. Pathways enriched in differentially variable positions identified in RA discordant twins, restricted to sites which were hypervariable in healthy co-twins. Pathway analysis was performed using the gometh function in the MissMethyl package. Pathways are ranked by P-value (P<0.05).

## REFERENCES

1. Silman AJ, MacGregor AJ, Thomson W, Holligan S, Carthy D, Farhan A, Ollier WE: Twin concordance rates for rheumatoid arthritis: results from a nationwide study. Br J Rheumatol 1993, 32:903–907.

2. Silman AJ, Newman J, MacGregor AJ: Cigarette smoking increases the risk of rheumatoid arthritis. Results from a nationwide study of disease-discordant twins. Arthritis Rheum 1996, 39:732–735.

3. Hoovestol RA, Mikuls TR: Environmental exposures and rheumatoid arthritis risk. Curr Rheumatol Rep 2011, 13:431–439.

4. Breitling LP, Yang R, Korn B, Burwinkel B, Brenner H: Tobacco-smoking-related differential DNA methylation: 27K discovery and replication. Am J Hum Genet 2011, 88:450–457.

5. van Steenbergen HW, Luijk R, Shoemaker R, Heijmans BT, Huizinga TW, van der Helm-van Mil AH: Differential methylation within the major histocompatibility complex region in rheumatoid arthritis: a replication study. Rheumatology (Oxford) 2014, 53:2317–2318.

6. Nakano K, Boyle DL, Firestein GS: Regulation of DNA methylation in rheumatoid arthritis synoviocytes. J Immunol 2013, 190:1297–1303.

7. Liu Y, Aryee MJ, Padyukov L, Fallin MD, Hesselberg E, Runarsson A, Reinius L, Acevedo N, Taub M, Ronninger M, et al: Epigenome-wide association data implicate DNA methylation as an intermediary of genetic risk in rheumatoid arthritis. Nat Biotechnol 2013, 31:142–147.

8. Glossop JR, Emes RD, Nixon NB, Haworth KE, Packham JC, Dawes PT, Fryer AA, Mattey DL, Farrell WE: Genome-wide DNA methylation profiling in rheumatoid arthritis identifies disease-associated methylation changes that are distinct to individual T- and B-lymphocyte populations. Epigenetics 2014, 9:1228–1237.

9. de la Rica L, Urquiza JM, Gomez-Cabrero D, Islam AB, Lopez-Bigas N, Tegner J, Toes RE, Ballestar E: Identification of novel markers in rheumatoid arthritis through integrated analysis of DNA methylation and microRNA expression. J Autoimmun 2013, 41:6–16.

10. Julia A, Absher D, Lopez-Lasanta M, Palau N, Pluma A, Waite Jones L, Glossop JR, Farrell WE, Myers RM, Marsal S: Epigenome-wide association study of rheumatoid arthritis identifies differentially methylated loci in B cells. Hum Mol Genet 2017, 26:2803–2811.

11. Bell JT, Spector TD: DNA methylation studies using twins: what are they telling us? Genome Biol 2012, 13:172.

12. Javierre BM, Fernandez AF, Richter J, Al-Shahrour F, Martin-Subero JI, Rodriguez-Ubreva J, Berdasco M, Fraga MF, O’Hanlon TP, Rider LG, et al: Changes in the pattern of DNA methylation associate with twin discordance in systemic lupus erythematosus. Genome Res 2010, 20:170–179.

13. Gomez-Cabrero D, Almgren M, Sjoholm LK, Hensvold AH, Ringh MV, Tryggvadottir R, Kere J, Scheynius A, Acevedo N, Reinius L, et al: High-specificity bioinformatics framework for epigenomic profiling of discordant twins reveals specific and shared markers for ACPA and ACPA-positive rheumatoid arthritis. Genome Med 2016, 8:124.

14. Tsai PC, Bell JT: Power and sample size estimation for epigenome-wide association scans to detect differential DNA methylation. Int J Epidemiol 2015.

15. Paul DS, Teschendorff AE, Dang MA, Lowe R, Hawa MI, Ecker S, Beyan H, Cunningham S, Fouts AR, Ramelius A, et al: Increased DNA methylation variability in type 1 diabetes across three immune effector cell types. Nat Commun 2016, 7:13555.

16. Teschendorff AE, Gao Y, Jones A, Ruebner M, Beckmann MW, Wachter DL, Fasching PA, Widschwendter M: DNA methylation outliers in normal breast tissue identify field defects that are enriched in cancer. Nat Commun 2016, 7:10478.

17. Teschendorff AE, Widschwendter M: Differential variability improves the identification of cancer risk markers in DNA methylation studies profiling precursor cancer lesions. Bioinformatics 2012, 28:1487–1494.

18. Hansen KD, Timp W, Bravo HC, Sabunciyan S, Langmead B, McDonald OG, Wen B, Wu H, Liu Y, Diep D, et al: Increased methylation variation in epigenetic domains across cancer types. Nat Genet 2011, 43:768–775.

19. Moayyeri A, Hammond CJ, Valdes AM, Spector TD: Cohort Profile: TwinsUK and healthy ageing twin study. Int J Epidemiol 2013, 42:76–85.

20. Aryee MJ, Jaffe AE, Corrada-Bravo H, Ladd-Acosta C, Feinberg AP, Hansen KD, Irizarry RA: Minfi: a flexible and comprehensive Bioconductor package for the analysis of Infinium DNA methylation microarrays. Bioinformatics 2014, 30:1363–1369.

21. Tian Y, Morris TJ, Webster AP, Yang Z, Beck S, Feber A, Teschendorff AE: ChAMP: Updated Methylation Analysis Pipeline for Illumina BeadChips. Bioinformatics 2017.

22. Barfield RT, Kilaru V, Smith AK, Conneely KN: CpGassoc: an R function for analysis of DNA methylation microarray data. Bioinformatics 2012, 28:1280–1281.

23. Reinius LE, Acevedo N, Joerink M, Pershagen G, Dahlen SE, Greco D, Soderhall C, Scheynius A, Kere J: Differential DNA methylation in purified human blood cells: implications for cell lineage and studies on disease susceptibility. PLoS One 2012, 7:e41361.

24. Houseman EA, Accomando WP, Koestler DC, Christensen BC, Marsit CJ, Nelson HH, Wiencke JK, Kelsey KT: DNA methylation arrays as surrogate measures of cell mixture distribution. BMC Bioinformatics 2012, 13:86.

25. Benjamini Y, Hochberg Y: Controlling the False Discovery Rate - a Practical and Powerful Approach to Multiple Testing. Journal of the Royal Statistical Society Series B-Methodological 1995, 57:289–300.

26. Bonder MJ, Luijk R, Zhernakova DV, Moed M, Deelen P, Vermaat M, van Iterson M, van Dijk F, van Galen M, Bot J, et al: Disease variants alter transcription factor levels and methylation of their binding sites. Nat Genet 2017, 49:131–138.

27. Guilhamon P, Eskandarpour M, Halai D, Wilson GA, Feber A, Teschendorff AE, Gomez V, Hergovich A, Tirabosco R, Fernanda Amary M, et al: Meta-analysis of IDH-mutant cancers identifies EBF1 as an interaction partner for TET2. Nat Commun 2013, 4:2166.

28. Phipson B, Maksimovic J, Oshlack A: missMethyl: an R package for analyzing data from Illumina’s HumanMethylation450 platform. Bioinformatics 2016, 32:286–288.

29. Geeleher P, Hartnett L, Egan LJ, Golden A, Raja Ali RA, Seoighe C: Gene-set analysis is severely biased when applied to genome-wide methylation data. Bioinformatics 2013, 29:1851–1857.

30. Subramanian A, Tamayo P, Mootha VK, Mukherjee S, Ebert BL, Gillette MA, Paulovich A, Pomeroy SL, Golub TR, Lander ES, Mesirov JP: Gene set enrichment analysis: a knowledge-based approach for interpreting genome-wide expression profiles. Proc Natl Acad Sci U S A 2005, 102:15545–15550.

31. Okada Y, Wu D, Trynka G, Raj T, Terao C, Ikari K, Kochi Y, Ohmura K, Suzuki A, Yoshida S, et al: Genetics of rheumatoid arthritis contributes to biology and drug discovery. Nature 2014, 506:376–381.

32. Lotem J, Levanon D, Negreanu V, Bauer O, Hantisteanu S, Dicken J, Groner Y: Runx3 at the interface of immunity, inflammation and cancer. Biochim Biophys Acta 2015, 1855:131–143.

33. Tokuhiro S, Yamada R, Chang X, Suzuki A, Kochi Y, Sawada T, Suzuki M, Nagasaki M, Ohtsuki M, Ono M, et al: An intronic SNP in a RUNX1 binding site of SLC22A4, encoding an organic cation transporter, is associated with rheumatoid arthritis. Nat Genet 2003, 35:341–348.

34. Klunker S, Chong MM, Mantel PY, Palomares O, Bassin C, Ziegler M, Ruckert B, Meiler F, Akdis M, Littman DR, Akdis CA: Transcription factors RUNX1 and RUNX3 in the induction and suppressive function of Foxp3+ inducible regulatory T cells. J Exp Med 2009, 206:2701–2715.

35. Silva GM, Finley D, Vogel C: K63 polyubiquitination is a new modulator of the oxidative stress response. Nat Struct Mol Biol 2015, 22:116–123.

36. Quinonez-Flores CM, Gonzalez-Chavez SA, Del Rio Najera D, Pacheco-Tena C: Oxidative Stress Relevance in the Pathogenesis of the Rheumatoid Arthritis: A Systematic Review. Biomed Res Int 2016, 2016:6097417.

37. Plant D, Webster A, Nair N, Oliver J, Smith SL, Eyre S, Hyrich KL, Wilson AG, Morgan AW, Isaacs JD, et al: Differential Methylation as a Biomarker of Response to Etanercept in Patients With Rheumatoid Arthritis. Arthritis Rheumatol 2016, 68:1353–1360.

38. Schett G, Tohidast-Akrad M, Smolen JS, Schmid BJ, Steiner CW, Bitzan P, Zenz P, Redlich K, Xu Q, Steiner G: Activation, differential localization, and regulation of the stress-activated protein kinases, extracellular signal-regulated kinase, c-JUN N-terminal kinase, and p38 mitogen-activated protein kinase, in synovial tissue and cells in rheumatoid arthritis. Arthritis Rheum 2000, 43:2501–2512.

39. Maurice MM, Nakamura H, Gringhuis S, Okamoto T, Yoshida S, Kullmann F, Lechner S, van der Voort EA, Leow A, Versendaal J, et al: Expression of the thioredoxin-thioredoxin reductase system in the inflamed joints of patients with rheumatoid arthritis. Arthritis Rheum 1999, 42:2430–2439.

40. Yoshida S, Katoh T, Tetsuka T, Uno K, Matsui N, Okamoto T: Involvement of thioredoxin in rheumatoid arthritis: its costimulatory roles in the TNF-alpha-induced production of IL-6 and IL-8 from cultured synovial fibroblasts. J Immunol 1999, 163:351–358.

41. Johnstone SE, Baylin SB: Stress and the epigenetic landscape: a link to the pathobiology of human diseases? Nat Rev Genet 2010, 11:806–812.

42. Nakano K, Whitaker JW, Boyle DL, Wang W, Firestein GS: DNA methylome signature in rheumatoid arthritis. Ann Rheum Dis 2013, 72:110–117.

43. Chen Z, Miao F, Paterson AD, Lachin JM, Zhang L, Schones DE, Wu X, Wang J, Tompkins JD, Genuth S, et al: Epigenomic profiling reveals an association between persistence of DNA methylation and metabolic memory in the DCCT/EDIC type 1 diabetes cohort. Proc Natl Acad Sci U S A 2016, 113:E3002–3011.

44. Ziller MJ, Gu H, Muller F, Donaghey J, Tsai LT, Kohlbacher O, De Jager PL, Rosen ED, Bennett DA, Bernstein BE, et al: Charting a dynamic DNA methylation landscape of the human genome. Nature 2013, 500:477–481.

